# Metal fluorides - multi-functional tools for the study of phosphoryl transfer enzymes

**DOI:** 10.1101/2024.03.25.586559

**Authors:** Erika Pellegrini, Pauline Juyoux, Jill von Velsen, Nicola J. Baxter, Hugh R. W. Dannatt, Jin Yi, Matthew J. Cliff, Jonathan P. Waltho, Matthew W. Bowler

## Abstract

Enzymes facilitating the transfer of phosphate groups constitute the most extensive protein group across all kingdoms of life, making up approximately 10% of the proteins found in the human genome. Understanding the mechanisms by which enzymes catalyse these reactions is essential in characterising the processes they regulate. Metal fluorides can be used as multifunctional tools for the study of these enzymes. These ionic species bear the same charge as phosphate and the transferring phosphoryl group and, in addition, allow the enzyme to be trapped in catalytically important states with spectroscopically sensitive atoms interacting directly with active site residues. The ionic nature of these phosphate surrogates also allows their removal and replacement with other analogues. Here, we describe the best practices to obtain these complexes, their use in NMR, X-ray crystallography, cryoEM and SAXS and describe a new metal fluoride, scandium tetrafluoride, which has significant anomalous signal with soft X-rays.

**Highlights:**

⍰ Enzymes that catalyse phosphoryl transfer are the largest family of enzymes and are involved in the storage and transmission of genetic information, energy transfer, signalling and cellular differentiation
⍰ Metal fluorides form a comprehensive tool kit to study the mechanisms of these enzymes by stabilizing the active conformation; mimicking both the transition state and ground state; and placing spectroscopically sensitive atoms into the active site
⍰ A guide is presented to the optimal formation of these complexes and their use in a wide variety of techniques in structural biology

## Introduction and Historical Perspective

Enzymes that catalyse phosphoryl transfer are the largest superfamily of enzymes and have roles in numerous crucial cellular functions. Understanding the methods that have evolved to enable the movement of phosphoryl groups is necessary to gain insight into the many processes that it controls^1,2^. Metal fluoride moieties can occupy the active sites of phosphoryl transfer enzymes at the location of the transferring phosphoryl group and have been used to study these proteins by X-ray crystallography and NMR with excellent results. More recently, new metal fluoride complexes have been defined and characterized. Additionally, various other applications for these complexes have been discovered, expanding the toolkit accessible to scientists studying phosphoryl transfer enzymes.

Fluoride anions have been known for more than 50 years to modulate the activity of a range of phosphoryl transfer enzymes such as kinases, phosphorylases and phosphatases. However, the precise mechanism remained unknown until the discovery that millimolar concentrations of fluoride can leach aluminium cations from laboratory glassware. Aluminium and fluoride together in the presence of GDP were discovered to stimulate the activity of small G proteins^3^. Upon the solution of the structure of transducin α in 1994^4^, the nature of the species was confirmed as AlF_4_^−^, where it was shown to mimic the transition state of phosphoryl transfer. In the crystal structure, the octahedral AlF_4_ ^−^ moiety was located in the site usually occupied by the γ-phosphate of GTPγS and coordinated axially by the oxygen atoms of GDP (β-oxygen) and an ordered water molecule positioned for in-line nucleophilic attack.

Although AlF_4_ ^−^ is isoelectronic with the transferring phosphoryl group, the octahedral geometry places demands on the active site architecture evolved to coordinate a trigonal bipyramidal transition state. However, this elegant use of a metal-fluoride moiety mimicking the γ-phosphate geometry of GTP opened the possibility to ‘trap’ transition state analogue (TSA) complexes of other phosphoryl transfer enzymes, which led to a number of seminal structures that defined the important features needed for catalysis^5–9^.

Aluminium fluoride has since continued to play a leading role in the determination of important high-resolution TSA structures. At the turn of the century a new metal fluoride species was discovered that self-assembled in the active site and comprised a magnesium cation as the central metal, coordinated axially by donor and acceptor oxygen atoms, together with three fluoride atoms arranged equatorially^10^. With this trigonal bipyramidal geometry the MgF_3_ ^−^ moiety thereby represents a closer isoelectronic and isosteric mimic to the transition state of the transferring phosphoryl group than AlF_4_^−^.

Early metal fluoride screening experiments^11^ identified that beryllium trifluoride also possesses enzyme activation and inhibitory properties. However, unlike aluminium fluoride and magnesium fluoride, species which can adopt near-transition state geometries, the coordination of beryllium fluoride is strictly tetrahedral in enzyme complexes and thereby is an excellent mimic of a ground state phosphate group. A wide variety of enzyme complexes containing a BeF_3_ ^−^ species have been described crystallographically^12–16^.

## Comparison of metal fluorides and phosphate groups

The three primary metal fluorides described above provide the perfect tool kit to study phosphoryl transfer enzymes. Fluorine and oxygen have very similar atomic radii (Figure 1A), share the same valence orbitals and have strong propensities to form hydrogen bonds with hydrogen donors, making fluorine an excellent surrogate for oxygen. The three metals are close to phosphorus both in atomic size and periodicity but each has different properties, and different electrons available for hybridization, leading to different possible coordination numbers and geometries (Figures 1B and 2A). Together with fluoride, these metals form excellent analogues of phosphoryl groups in the active sites of enzymes.

**Figure 1.**
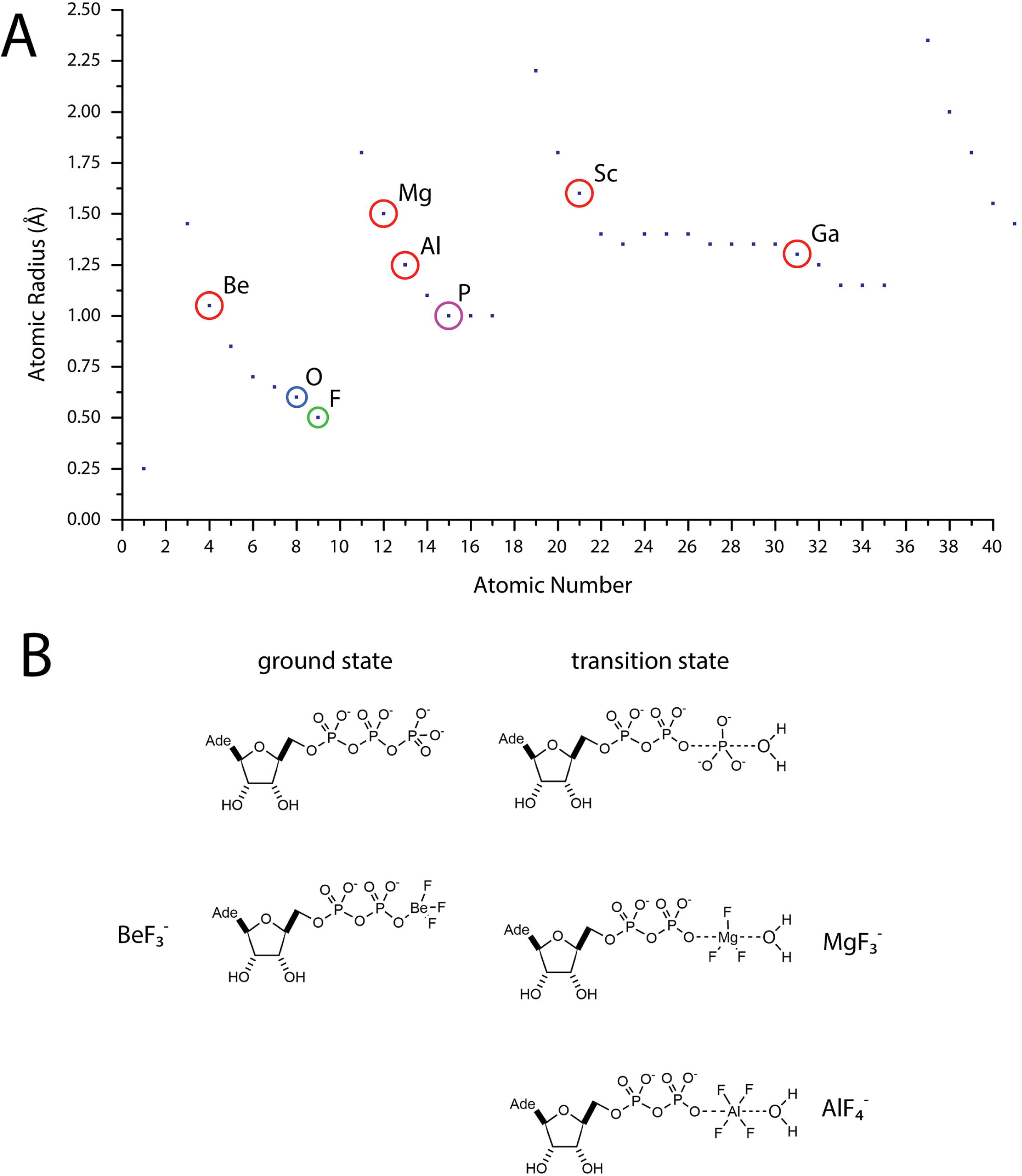
Metal fluorides. **A**. Atomic radii of the elements. The atomic radii (empirically measured covalent radii) of the elements are plotted in Ångstroms against the atomic number. The close relationship between phosphorus and the metals that form phosphoryl analogues with fluoride is shown. The scandium-fluoride bond is the longest observed at 2 Å and probably represents the limit for the largest atomic radius tolerated in the active sites of phosphoryl transfer enzymes. **B**. Schematic of metal fluorides compared to the ground state and transition state of the hydrolysis of ATP as an example.

**Figure 2.**
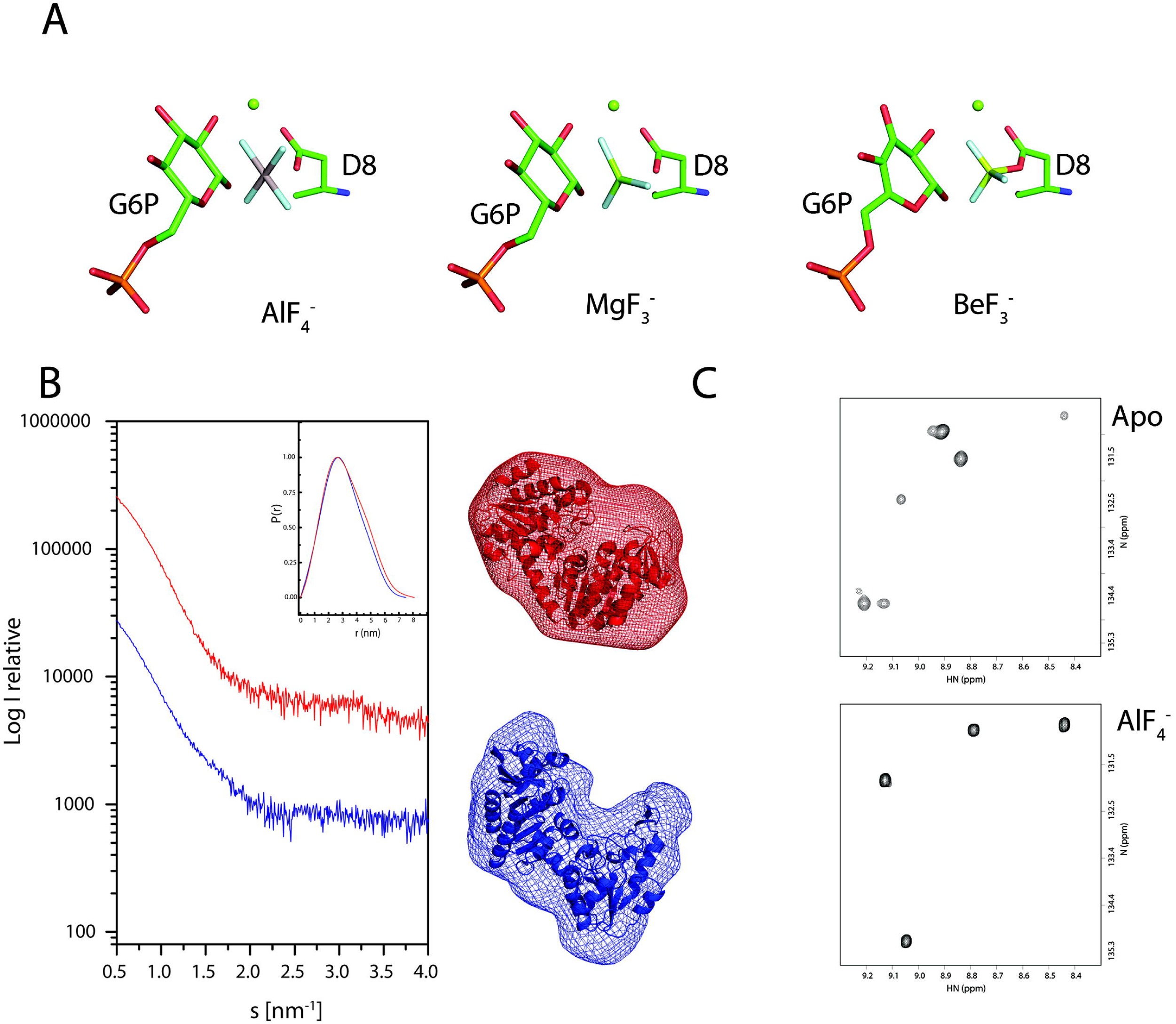
Metal fluorides. **A**. Examples of enzyme-metallofluoride complexes. AlF_4_^−^ and MgF_3_^−^ are shown in the active site of βPGM where they mimic the transition state of the phosphoryl transfer from glucose-1,6-bisphosphate (G6P) from the catalytic aspartate (D8). BeF_3_^−^ is shown in the active site of βPGM where it mimics the phosphoaspartyl intermediate of the catalytic cycle. **B**. Aluminium fluoride traps the active state in solution allowing it to be compared to the *apo* enzyme. Scattering curves, distance distribution functions (inset chart) and *ab initio* shapes determined from SAXS data recorded on *apo* PGK and the PGK-3PG-AlF_4_^−^-3PG TSA complex and the structures fit to the envelopes. Data from the *apo* protein are shown in *blue* and from the aluminium fluoride complex *red*. **C.** Identical regions of ^1^H-^15^N TROSY spectra recorded on substrate free βPGM and the βPGM-AlF_4_^−^-G6P TSA complexes using standard NMR conditions. For substrate free βPGM slow exchange dynamics result in the presence of multiple peaks for selected residues. For the βPGM-AlF_4_^−^-G6P TSA complex, single peaks indicate that these slow dynamics have been arrested confirming the ability of AlF_4_^−^ to act as a molecular clamp.

## Principal advantages of the use of metal fluorides

### Trap enzymes in stable, functionally-relevant conformations

Many enzymes perform catalysis by involving large conformational changes throughout the reaction coordinate such as the re-structuring of mobile loops, domain-domain closure events or in the binding of large substrates such as RNA^6,17,18^. Such dynamics can make it difficult to either study catalytically relevant conformations or maintain a sufficiently homogenous population to allow crystallization to occur or enough particles to be selected to obtain a meaningful 3D reconstruction in cryogenic electron microscopy (cryoEM). By the formation of very stable complexes, the use of metal fluorides can remove or reduce protein flexibility and fix phosphate-transferring enzymes into an active and immobile state. Such states provide structural homogeneity and therefore excellent starting points for crystallization trials or other studies^19^.

### Study of the reaction mechanism; insights into catalysis

In addition to the structural possibilities provided by the use of transition state analogues, the ability to sample a ground-state analogue complex by the use of BeF_3_ ^−^ allows for comparison of structure throughout the various stages in the reaction coordinate. By combining a series of enzyme complexes containing metal fluoride analogues of the ground state and transition state, the interaction of active site residues with surrogates of the oxygen atoms of the phosphoryl group may be visualized for the complete phosphoryl transfer reaction.

### Convenience

Finally, metal fluoride complexes are both simple and inexpensive to prepare, and have potential application in any enzyme system that involves phosphoryl transfer. As they are intrinsically unreactive species, their use avoids the necessity of non-hydrolysable substrate analogues that often have lower binding affinities than the cognate substrate, and may introduce structural distortions. The presence of electron-rich metal ions and fluorine nuclei in the active site also opens specific avenues for X-ray diffraction and NMR analyses.

Here, we describe the best protocols to obtain these complexes, their various uses in structural biology and present the structure of a new metal fluoride, scandium tetrafluoride, that places an anomalously scattering element in the active site of phosphoryl transfer enzymes and compare it with the equivalent AlF_4_^−^ complex.

## Choosing the right tool for the job

### AlF_4_^−^

The most commonly used metal fluoride ion, aluminium tetrafluoride, has octahedral geometry (Figures 1B and 2A) and forms the most stable complexes with proteins and substrates. Its charge density is the highest and, as it is also present as a free species in solution, it does not require a fully formed active site in which to assemble. This makes it the first choice as a ‘molecular clamp’ to trap the active state of an enzyme in solution, despite having geometry different from that of the native transition state. We have shown that AlF_4_^−^ stabilizes these conformations in solution before crystallization using both SAXS and NMR (Figure 2B and C) thereby trapping the conformation for further study. AlF_4_^−^ stabilizes these states by strongly coordinating substrates and active site residues.

### MgF_3_^−^

Magnesium trifluoride ions also form transition state analogue complexes, but these have different properties from those formed by AlF_4_^−^. First, as well as being isoelectronic with the transferring phosphoryl group, MgF_3_^−^ ions are also isosteric, adopting a trigonal bipyramidal geometry that mimics the transition state moiety (Figures 1B and 2A). The combination of these properties make trifluoromagnesate ions much closer analogues of the transition state of phosphoryl transfer. However, the formation constant of MgF_3_^−^ ions in free solution is extremely low, which demands that the ion can only assemble directly in the active sites of enzymes^20^. As a result, the overall affinity of the MgF_3_^−^ ion is frequently found to be much lower than that of AlF_4_^−^. However, due to its closer match to the transition state, MgF_3_^−^ complexes are preferable for detailed analysis of catalytic mechanisms, compared to the ones formed by AlF_4_^−^.

### BeF_3_^−^

Beryllium trifluoride ions have obligate tetrahedral geometry and therefore form analogues that are isosteric and isoelectronic with the ground state of phosphoryl transfer (Figures 1B and 2A). The crystal structures of BeF_3_^−^ and MgF_3_ ^−^ complexes place the fluoride ions in the locations of both ground and transition states and therefore provide the best starting points for modeling studies of the enzyme active site. BeF_3_^−^ also has a role for proteins other than phosphoryl transfer enzymes, as it has been successfully used to mimic the phosphate moiety bound at phosphorylation sites, leading to structural and functional properties which mimic those of the phosphorylated protein^13,21^. This behavior is particularly useful as homogenous populations of phosphoproteins can be difficult to prepare.

## Forming metal fluoride complexes

The model systems β-phosphoglucomutase (βPGM) and human phosphoglycerate kinase (PGK) catalyse different phosphoryl transfer reactions and in combination with metal fluoride moieties both ground state and transition state analogue complexes have been investigated using a variety of structural techniques to define several aspects of catalysis. Here, we use these systems to illustrate how to form the complexes and how to best exploit their various properties.

The great advantage of metal fluoride TSA and ground state analogue (GSA) complexes is that all components are present in solution and readily self-assemble in the active site forming stable enzyme complexes that are relevant to the catalytic cycle. The inorganic metal fluoride salts (AlF_3_, and MgF_2_) are too insoluble to use and therefore, the fluoride anion and metal cation components must be added from separate stock solutions. Both ammonium fluoride and sodium fluoride are suitable as the source of fluoride and are readily soluble in water. No difference is observed in the formation of the metal fluoride protein complex but a difference may be apparent in the crystallization trials where the counter ion can have an effect on crystal packing. Metal chlorides, such as AlCl_3_, MgCl_2_, BeCl_2_, GaCl_3_ and ScCl_3_ can be easily dissolved in water at high concentration (0.5 M) and the solutions conserved at −20°C. One of the critical aspects in preparing metal fluoride enzyme complexes is the pH of the resulting solution. In particular, solutions of AlCl_3_, BeCl_2_, GaCl_3_ and ScCl_3_ are highly acidic (pH ∼2) and the addition of even small volumes of a solution at this pH value to a protein sample will cause precipitation. As the tolerance of protein precipitation within samples is quite different for crystallization and NMR studies, the preparation process of the metal chloride stock solutions and the protein samples is optimized in different ways. Of particular importance is the pH adjustment of the stock solutions and the sequence of component addition.

In transition state analogue complexes, the axial ligands of the metal fluoride ions correspond to the phosphate donor and acceptor oxygen atoms of the transferring phosphate group. Therefore, for the formation of the complex the native substrate(s) – lacking the phosphate which the metal fluoride mimics – must also be present in solution. For example: β-PGM requires glucose-6-phosphate (G6P) or β-glucose-1-phosphate (Figure 2A), phosphoserine phosphatase requires serine or H_2_O and phosphoglycerate kinase requires both ADP and 3-phosphoglycerate (3PG). In most cases, such metal fluoride complexes require the presence of the product or cofactor of the enzymatic reaction in stoichiometric excess.

BeF_3_^−^ ground state complexes have been shown to form either when both phosphate acceptor and donor molecules are present, or when only one is present. For example, BeF_3_^−^ complexes of β-PGM can form in the presence or absence of the phosphate acceptor molecule G6P. In both cases, the BeF_3_^−^ ion is found to be axially coordinated by the catalytic aspartate residue (D8) that acts as the nucleophile in the active site of β-PGM (Figure 2A). Indeed in all BeF_3_^−^ complexes of phosphoryl transfer enzymes studied thus far, the BeF_3_ ^−^ ion is found coordinated to the most acidic of the two available acceptor/donor groups^16^.

### Sample preparation for crystallization, SAXS and cryoEM

In order to increase the pH values of the metal chloride stock solutions, they should be prepared in 100 mM unbuffered Tris base. In the case where an AlF_4_^−^ complex is desired, the protein buffer must be maintained below pH 7.5 to guarantee the aluminium cation solubility. Titration experiments monitored using ^19^F NMR spectroscopy have shown that AlF_4_ ^−^ and AlF_3_(OH)^−^ species in solution are pH dependent and above pH 8 aluminium cations are precipitated from solution as insoluble aluminium hydroxide^20,22^. Under these aluminium limited conditions, only trigonal bipyramidal MgF_3_^−^ TSA complexes can be obtained^16^. Therefore, the optimized sequence of component addition is to add fluoride to the prepared buffer first, then the metal chloride stock, thereby allowing the formation of metal fluoride solution species first. Following pH re-adjustment, the protein stock solution is added to the final buffer together with the substrate required for the TSA or GSA complexes. In this manner, the chances of forming stable metal fluoride complexes whilst maintaining solubility of both protein and metal cations are increased. Typical concentrations for the components are: 10-40 mM fluoride ions, 5-20 mM magnesium ions, 1-5 mM aluminium ions, 1-10 mM beryllium, 10 mM scandium and 10 mM gallium ions. These concentrations need to be reduced for cryoEM studies as high concentrations can increase background noise when the sample is small. In these cases, reducing the concentrations to 250 μM fluoride ions and 25 μM metal ions improves signal to noise significantly while maintaining the complex^23^.

### Sample preparation for NMR studies

The protein concentrations used in NMR samples are much higher than those used in crystallization experiments. Therefore, in order to avoid unnecessary dilution of the protein component (0.5-1.0 mM), the ingredients are added directly to the buffered protein stock at neutral pH containing an appropriate concentration of MgCl_2_ as the source of the metal catalytic cofactor and 1-2 mM NaN_3_ as a preservative. The metal chloride stock solution (AlCl_3_, BeCl_2_, ScCl_3_ or GaCl_3_) and fluoride stock solution (NH_4_F or NaF) are added first to inhibit any trace of residual enzyme activity, followed by the substrate. When sodium fluoride is used as the fluoride source, the buffering capacity of the protein sample against the basicity of NaF should be considered. Some kinases have less tolerance to the high concentration of NH_4_F present when using a protein buffer with a relatively neutral pH value. Any unwanted protein precipitate generated during the sample preparation procedure can be removed effectively by centrifugation prior to transferring the sample into the NMR tube. Typical concentrations for the components are: 10-40 mM fluoride ions, 5-20 mM magnesium ions, 1-3 mM aluminium ions, 1-5 mM beryllium ions, 10 mM scandium ions and 10 mM gallium ions. However, the fluoride concentrations used here can leach aluminium ions from glass NMR tubes, (Wilmad-LabGlass, NJ) particularly if in contact for extended periods (>2-3 days), and so for experiments involving MgF_3_^−^ TSA complexes, the sample should be supplemented with 1-2 mM deferoxamine, which is a potent chelator of aluminium cations. For ^19^F NMR experiments recorded in 100% H_2_O buffer, the deuterium NMR lock is provided by external 100% D_2_O buffer present in a sealed glass capillary tube inserted in the conventional NMR tube.

## X-Ray Crystallography Methods

### Cross-soaking

The ability to trap proteins with a TSA can help in the understanding of the reaction mechanism but reveals little information concerning the binding of the ground states in the reaction coordinate, or of the binding of other ligands or inhibitors. Attempts to co-crystallise the closed conformation of PGK in the presence of ADP, ATP analogues (AMP-PNP and AMP-PCP) or BeF_3_ ^−^ failed as these complexes cannot maintain the closed state, presenting a flexible complex for crystallization^19^.We have found that for PGK, MgF_3_^−^ and AlF_4_^−^ can be used to trap the closed conformation for crystallization and subsequently be removed and a new ligand soaked in while maintaining the closed conformation, a practice known as cross-soaking that has been shown to be effective with peptides and other ligands^24–27^. As metal fluorides are ionic species, this allows the ions to diffuse out from the active site when the crystals are presented with a solution where they are absent. The lower intrinsic stability of MgF_3_^−^ means that it can be diffuse out the active site when placed in a buffer lacking fluoride ions, but we have successfully removed both aluminium and magnesium species. Crystal packing maintains the conformation initially stabilized by the metal fluoride and allows alternative ligands to be inserted – in this way the active conformation in the presence of an ATP analogue, the BeF_3_^−^ complex mimicking the 1,3-bisphosphoglycerate (1,3BPG) complex were determined (see Table 1 and supplementary information for detailed crystallographic methods). Crystals of the MgF_3_^−^, ADP and 3PG quaternary complex (PGK-3PG-MgF_3_^−^-ADP TSA-complex) were soaked in buffers that matched the mother liquor but were devoid of fluoride ions. Soaking times were varied and datasets collected on crystals harvested at 10 minute intervals (see Table 1 for details of data reduction and refinement and supplementary information for detailed crystallographic methods). The optimal time for complete removal of the metal fluoride was found to be 30 minutes (this will of course vary, depending on the system being studied, the solute accessibility of the active site and crystal packing) and the absence of the TSA was confirmed by analysis of the difference Fourier maps (Figure 3A). A further 30 minutes was then required to allow AMP-PCP to enter the active site and displace the remaining ADP (Figure 3B). The complex with the phosphonate analogue AMP-PCP was only obtained with the K219A varient (HsPGK(K219A)-3PG-AMPPCP GSA-complex). The protonation of the phosphonate is favorable in solution under the experimental conditions, and so it might be expected to readily form the inter-substrate hydrogen bond seen in TSA structures. However, the hydrogen bond network cannot accommodate the bridging methylene of AMP-PCP which could prevent binding. For the PGK-3PG-BeF_3_ ^−^-ADP GSA complex the above procedure was followed except that after removal of the metal fluoride the crystal was then placed into the mother liquor containing 10 mM ammonium fluoride and 20 mM BeCl_2_ for 30 minutes, to introduce the ground state analogue (Figure 3C). The high concentration of beryllium was needed to compete with any formation of MgF_3_^−^ from the catalytic magnesium present. Using cross-soaking TSAs were used to define the entire reaction coordinate of the protein from the ground state of 1,3BPG to ATP (Figure 3D).

**Figure 3.**
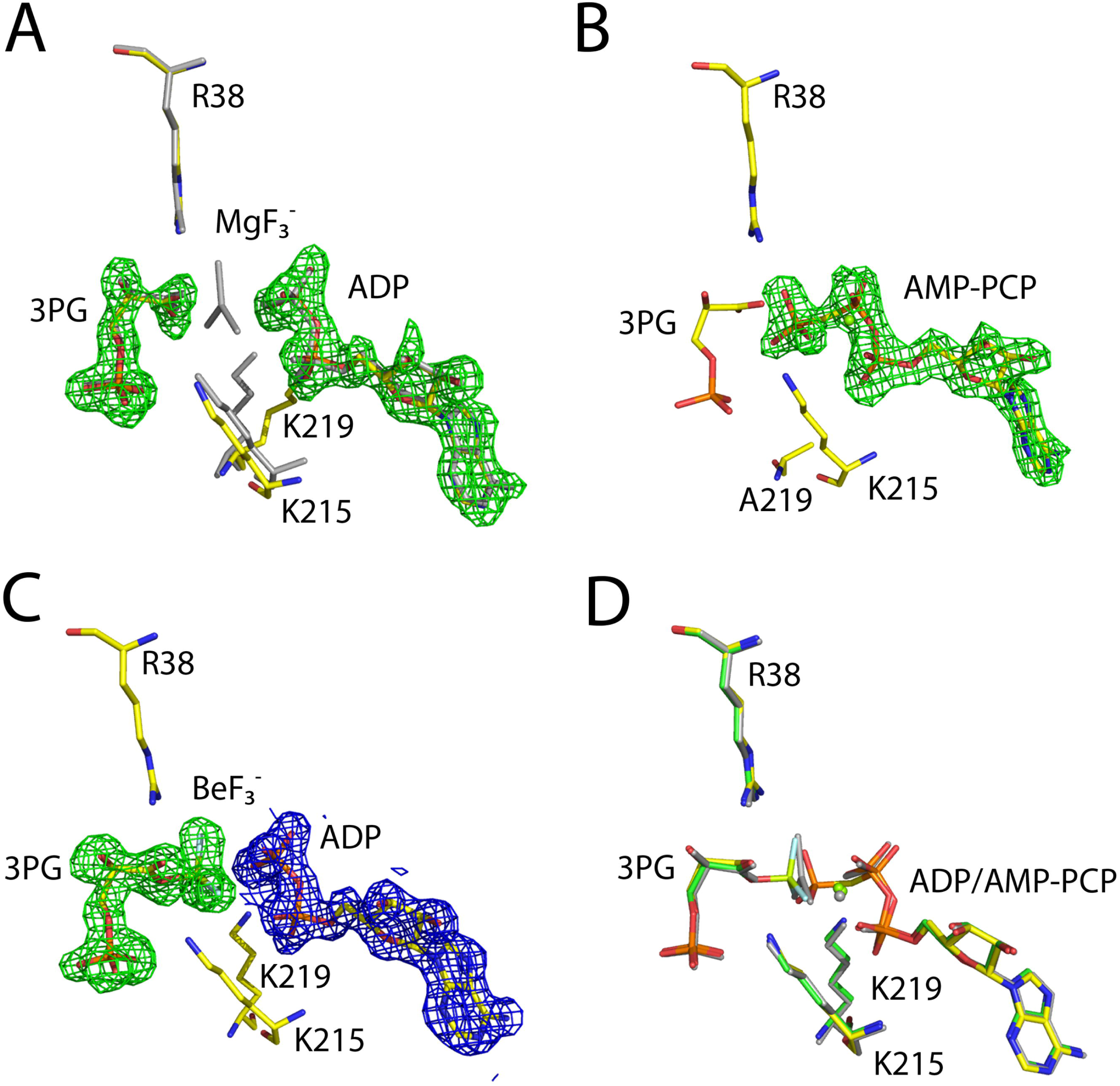
Cross soaking to obtain ground state structures of PGK. **A**. Initially crystals of PGK-3PG-MgF_3_^−^-ADP TSA complex were soaked in a buffer lacking fluoride ions and the structure determined. The difference Fourier demonstrates the complete removal of the transition state analogue and the movement of a catalytic lysine (K219) also confirms the removal (the PGK-3PG-MgF_3_^−^-3PG TSA complex is shown in gray for reference). **B**. Once the transition state analogue is removed new ligands can be soaked in – here, AMP-PCP replaces ADP showing the ATP ground state. **C**. When beryllium ions are added the map shows the ground state analogue coordinated to 3PG mimicking the 1,3-BPG complex. When combined with the transition state analogue complexes a complete description of catalysis is obtained **D**. Difference maps (green) are shown contoured at 3σ and 2mF_o_−F_c_ maps (blue) are shown contoured at 1.5σ.

**Table 1.**
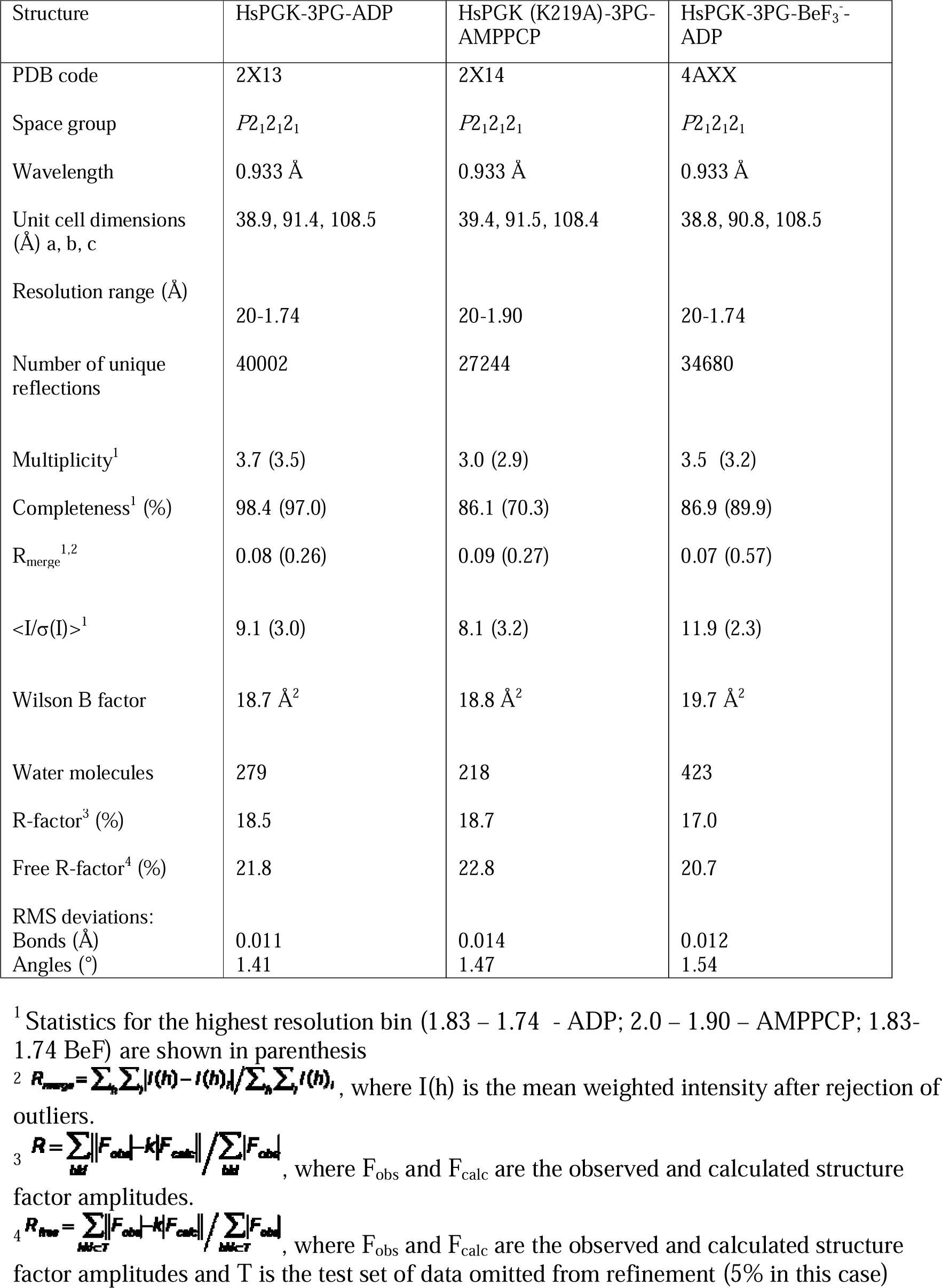
Data collection and refinement statistics for PGK closed conformations and phosphate identification.

### Potential phasing applications of metal fluoride complexes

A number of derivatised nucleotide compounds have been developed that allow the insertion of a heavy atom into the active site of a nucleotide binding protein. These nucleotides contain a halogen^27,28^ or a selenium atom^29^ that serve as anomalous scattering centres, either to locate the nucleotide binding site in low resolution structures^30^ or to be used in MAD or SAD phasing experiments^31^. These compounds are enormously useful; however, they are only effective for proteins that bind nucleotides and can be costly to synthesise. Given that magnesium, aluminum and beryllium all form protein metal fluoride complexes, the possibility of making complexes with metals that have a K-absorption edge within the energy range available at synchrotron radiation sources can be investigated as a potential source of new phasing compounds. It has been reported that myosin can be inhibited by metal fluorides other than those containing aluminium, magnesium and beryllium^32^ and this was used as a starting point to screen, using ^19^F-NMR and crystallography, the presence of fluoride complexes with metals having an K-absorption edge relevant to protein crystallography. Using βPGM as a model, a number of metals were screened for the formation of protein-bound metal fluoride complexes and both scandium and gallium form tetrafluoride complexes were observed (Figure 4A). The structure of the scandium tetrafluoride complex of β-PGM (βPGM-ScF_4_ˉ-G6P TSA complex) was determined at 1.3 Å and compared to the aluminium complex (βPGM-AlF_4_ˉ-G6P TSA complex) determined at the same resolution; see Table 2 for details of data reduction and refinement (see supplementary information for detailed ^19^F-NMR and crystallographic methods).

**Figure 4.**
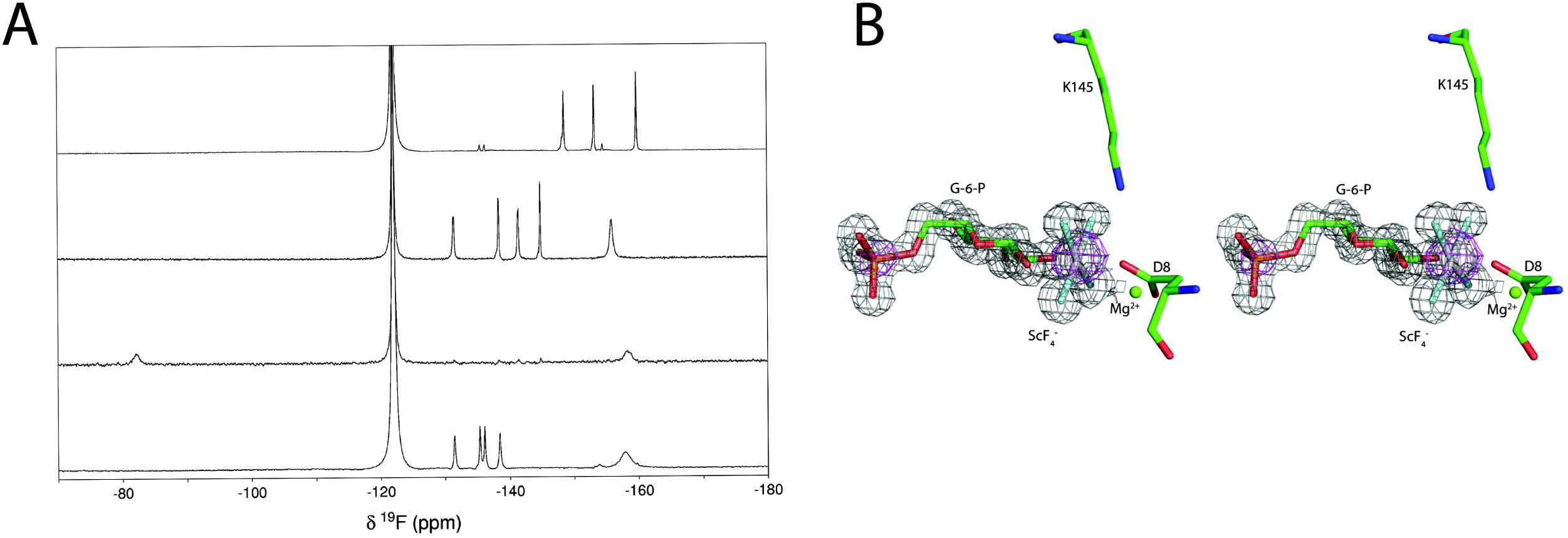
New metallofluorides. **A**. ^19^F NMR spectra for βPGM-G6P TSA complexes with MgF_3_^−^, AlF_4_^−^, ScF_4_^−^ and GaF_4_^−^ (top to bottom) occupying the site of the transferring phosphoryl group. The sharp GaF_4_^−^ complex resonances appear at similar frequencies to those of the previously reported AlF_4_^−^ complex, with chemical shifts of −131.4, −135.3, −136.1 and −138.4 ppm. In the ScF_4_^−^ complex, a single broad protein-dependent peak appears at −82.2 ppm, suggesting rotational averaging of the ^19^F environments in solution. In all cases the free F^−^ ion resonance appears at −122.0 ppm, and for the AlF_4_^−^, ScF_4_^−^ and GaF_4_^−^ complex spectra, the broad peaks upfield of −150 ppm are the result of free metal fluoride species in solution. Both Sc and Ga have K edges accessible to, or near, the energies available at synchrotron beamlines. **B**. Stereo image of the active site of the βPGM-ScF_4_^−^-G6P – structure. The 2mF_o_−F_c_ map from native data collected to 1.3 Å is shown as a grey mesh contoured at 1.5σ and the anomalous difference Fourier calculated from data collected at 7 keV is shown as a magenta mesh contoured at 5σ. The scandium atom is clearly identified by a 15.4σ peak in the anomalous difference maps; the corresponding peak for the phosphorus atom is 8.5σ.

**Table 2.** Data collection and refinement statistics for PGM structures.

| Structure | PGM-AlF<sub>4</sub><sup>-</sup>-G6P<br>native | PGM-ScF<sub>4</sub><sup>-</sup>-G6P<br>native | βPGM-ScF<sub>4</sub><sup>-</sup>-G6P<br>long wavelength |
| --- | --- | --- | --- |
| PDB code | 2WF6 | 3ZI4 |  |
| Space group | *P*2<sub>1</sub>2<sub>1</sub>2<sub>1</sub> | *P*2<sub>1</sub>2<sub>1</sub>2<sub>1</sub> | *P*2<sub>1</sub>2<sub>1</sub>2<sub>1</sub> |
| Wavelength (Å) | 0.933 | 0.933 | 1.77 |
| Unit cell dimensions (Å) a, b, c | 38.0, 54.4, 104.9 | 37.2, 54.2, 104.6 | 37.2, 54.2, 104.5 |
| Resolution range (Å) | 20.0 -1.3 | 20-1.33 | 20-1.95 |
| Number of unique reflections | 50761 | 46171 | 15433 |
| Multiplicity<sup>1</sup> | 3.2 (3.1) | 3.4 (3.0) | 12.4 (12.5) |
| Completeness<sup>1</sup> (%) | 93.5 (86.9) | 94.0 (76.6) | 96.3 (92.8) |
| R<sub>merge</sub><sup>1,2</sup> | 0.05 (0.26) | 0.05 (0.45) | 0.10 (0.21) |
| <I/σ(I)><sup>1</sup> | 9.2 (2.9) | 18.7 (2.4) | 19.1 (10.8) |
| Wilson B factor (Å<sup>2</sup>) | 10.4 | 10.7 | 14.6 |
| Water molecules | 318 | 422 |  |
| R-factor<sup>3</sup> | 15.0 | 13.0 |  |
| Free R-factor<sup>4</sup> | 18.6 | 17.4 |  |
| RMS deviations: |  |  |  |
| Bonds (Å) | 0.01 | 0.01 |  |
| Angles (°) | 1.46 | 1.67 |  |

<sup>1</sup> Statistics for the highest resolution bin (PGM-AlF<sub>4</sub><sup>-</sup>-G6P native: 1.37 - 1.30 Å; PGM-ScF<sub>4</sub><sup>-</sup>-G6P native: 1.4 – 1.33 Å and PGM- ScF<sub>4</sub><sup>-</sup>-G6P long wavelength: 2.06 –1.95 Å) are shown in parenthesis
<sup>2</sup> $R_{\text{merge}} = \sum_h \sum_i |I(h) - \bar{I}(h)| / \sum_h \sum_i I(h)$ , where I(h) is the mean weighted intensity after rejection of outliers.
<sup>3</sup> $R = \sum_h |F_{\text{obs}} - k|F_{\text{calc}}| / \sum_h |F_{\text{obs}}|$ , where F<sub>obs</sub> and F<sub>calc</sub> are the observed and calculated structure factor amplitudes.
<sup>4</sup> $R_{\text{free}} = \sum_{h \in T} |F_{\text{obs}} - k|F_{\text{calc}}| / \sum_{h \in T} |F_{\text{obs}}|$ , where F<sub>obs</sub> and F<sub>calc</sub> are the observed and calculated structure factor amplitudes and T is the test set of data omitted from refinement (5% in this case).

As expected, scandium (III) forms an octahedrally coordinated tetrafluoride moiety between the catalytic aspartate residue (D8) and the 1-hydroxyl group of G6P in the βPGM active site resulting in a near-transition state complex (Figure 4B). This novel ScF_4_^−^ species binds in a similar manner to its AlF_4_^−^ counterpart but has longer metal fluoride bonds, 2.0 Å vs 1.8 Å, respectively (Figure S1) and axial oxygen distances at 2.0 Å and 2.2 Å. The scandium atom was positively identified as occupying the position of the central metal by the collection of a data set at an energy of 7 keV. At this wavelength the anomalous scattering length from scandium is 1.95 electrons (against 0.56 electrons for phosphorus and 0.72 electrons for sulphur). Inspection of the anomalous difference Fourier maps revealed a peak of 15.4σ at the position of the central metal (Figure 4B), thereby confirming for the first time the atomic identity of the central metal in a metallofluoride. Therefore, ScF_4_^−^ species can be used to trap enzymes in near-transition state conformations, additionally identifying the location of active sites and facilitating SAD phasing protocols where it would contribute considerably to the anomalous signals from native sulphur and/or phosphorous atoms. Enzyme TSA complexes involving GaF_4_^−^ species would be extremely useful for MAD phasing experiments as its K-absorption edge is at 10.37 keV (λ=1.20 Å) within the energy range available at macromolecular crystallography beamlines. We obtained crystals of the ScF_4_^−^ complex but not of the corresponding gallium complex, though the latter was detectable in solution (Figure 4A), which may be due to the requirement of a low pH being incompatible with the crystallization conditions for β-PGM.

The metals that have been shown to form enzyme-metallofluoride complexes all fall within a close group and periodic range (Figure 1A). The ability of the metal fluoride moieties to self-assemble in the active sites of phosphoryl transfer enzymes is probably dependent on the electronic configuration but also the atomic radius of the atom. The metal-fluoride bonds that have been characterised vary from 1.6 Å in BeF_3_^−^ ^21^, 1.8 Å in AlF_4_^−^, 1.9 Å in MgF_3_^−^ ^10,33^ and 2.0 Å in ScF_4_^−^, whereas the phosphoryl oxygen bond varies between 1.5 and 1.7 Å. Therefore, the atomic radius of scandium (1.6 Å) probably represents the maximum for a metal fluoride that is readily tolerated in an enzyme active site.

## ^19^F-NMR

### ^19^F-NMR properties

^19^F NMR studies of protein complexes involving fluorinated compounds or metal fluoride moieties are attractive as the ^19^F nucleus occurs at 100% natural abundance, is spin ½ with high sensitivity (83% of ^1^H) and has a large chemical shift range (100-fold larger than ^1^H). The resulting spectra are also relatively simple to interpret and which are free from background signals^34,35^. Solution ^19^F NMR was first applied to protein systems in the late 1960s^36^ and one of the primary advantages of this technique is to report on the structures of known protein systems in their native solution environment. Since the ^19^F spectra for such complexes are relatively simple, the molecular weight restrictions that impact NMR approaches using multidimensional strategies (^1^H, ^13^C, ^15^N) are elevated such that systems as large as 200 kDa can be studied. The ^19^F chemical shift is extremely sensitive to changes in local electrostatics and alterations in van der Waals contacts. Hence, for enzyme-metallofluoride TSA and GSA complexes, ^19^F NMR spectroscopy offers an extremely sensitive approach with which to probe atomic environments in the vicinity of the phosphate oxygen atoms in catalytically relevant active site geometries. ^19^F chemical shift values, solvent induced isotope shifts (SIIS) observed on the ^19^F resonances, linewidth measurements, cross hydrogen bond F—HN scalar ^1^*J*_HF_ and ^2^*J*_NF_ couplings and driven-^201^H and transient-^201^H nuclear Overhauser effects (NOEs) all report on interatomic proximities in the solution state which can be correlated with high-resolution crystal structures.

### ^19^F chemical shifts

^19^F resonances display a very large chemical shift range (∼1000 ppm) and are predictable with good precision from quantum calculations of electronic distributions^37^. However, the chemical shift range observed for ^19^F resonances in enzyme-metallofluoride complexes is lower and in general they locate between −125 ppm and −195 ppm, when using CFCl_3_ resonating at 0.0 ppm as the IUPAC referencing standard. Since the ^19^F chemical shift is sensitive to the van der Waals environment as well as local electrostatic fields, the relative chemical shift positions within this range can reflect differential proton densities as shown by the significant upfield chemical shift observed for the most upfield peak between the βPGM-MgF_3_^−^ –G6P and βPGM-MgF_3_^−^ –2deoxyG6P TSA complexes (Figure 5A). Here, the removal of a contributing hydrogen bond between the sugar phosphate and the metal-fluoride results in the fluorine nucleus being significantly more shielded (Figure 5E).

**Figure 5.**
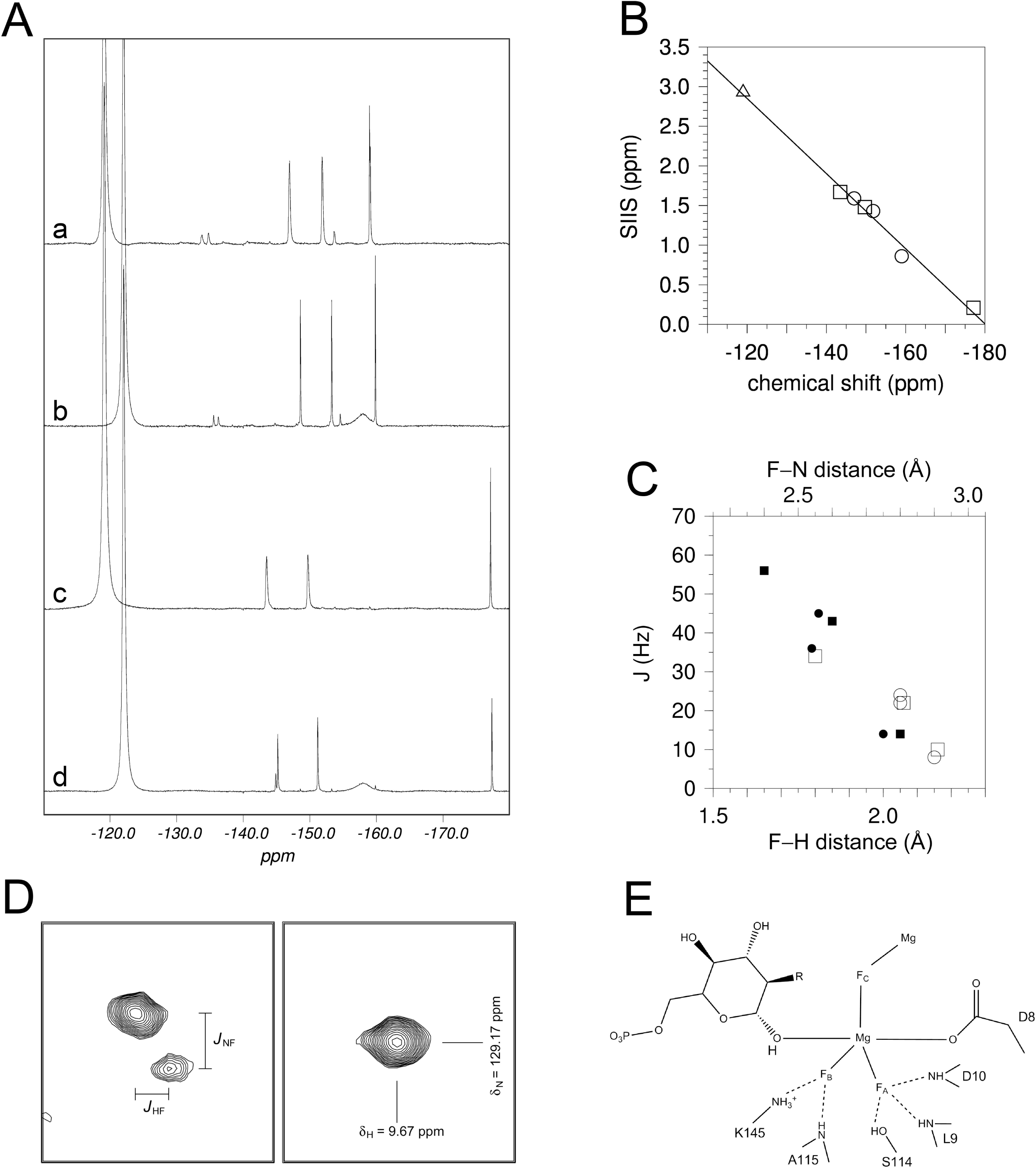
^19^F NMR parameters. **A**. ^19^F NMR spectra of βPGM-MgF_3_^−^ TSA complexes recorded at 25°C in either 50 mM HEPES 100% H_2_O or 100% D_2_O buffer, pH 7.2 with a) G6P in H_2_O buffer (F_A_ = −147.0, F_B_ = −151.8, F_C_ = −159.0 ppm), b) G6P in D_2_O buffer (F_A_ = −148.6, F_B_ = −153.3, F_C_ = −159.8 ppm), c) 2-deoxy-G6P in H_2_O buffer (F_A_ = −143.5, F_B_ = −149.7, F_C_ = −177.1 ppm) and d) 2-deoxy-G6P in D_2_O buffer (F = −145.2, F = −151.2, F = −177.3 ppm). Free F^−^ resonates at −119.0 ppm in 100% H_2_O buffer and at −122.0 ppm in 100% D_2_O buffer. **B.** Correlation plot showing the relationship between chemical shift (ppm) and SIIS (ppm) for the ^19^F resonance of free F^−^ (triangle) and those of the βPGM-MgF ^−^ –G6P TSA complex (circles) and the βPGM-MgF ^−^ –2-deoxy-G6P TSA complex (squares). Linear regression analysis gives R^2^ = 0.991. **C.** Correlation plot showing the relationships between *J*_HF_ (filled symbols) and *J*_NF_ (open symbols) couplings with the corresponding internuclear distances derived from structures of the βPGM-MgF ^−^ – G6P TSA (circles) and the βPGM-AlF ^−^ –G6P TSA (squares) complexes. The F-N distances are derived directly from the experimental coordinates and the F-H distances are determined to hydrogen atoms positioned using the program XPLOR. **D.** Identical regions of ^1^H, ^15^N HSQC spectra recorded without (left) and with (right) ^19^F decoupling showing the backbone amide resonance of A115 in the βPGM-MgF_3_^−^ G6P TSA complex. In the left spectrum, the HN peak is split horizontally by the ^1^H-^19^F coupling (*J*_HF_ = 36 Hz) and vertically by the ^15^N-^19^F coupling (*J*_NF_ = 24 Hz) through the hydrogen bond to F_B_ of the MgF moiety. In the right spectrum, these *J*-−coupling patterns collapse to an average position in a ^1^H, ^15^N HSQC spectrum recorded with ^19^F decoupling in both ^15^N and ^1^H dimensions. **E.** Schematic view of the βPGM-MgF_3_^−^ sugar phosphate TSA complex active site. In the βPGM-MgF_3_^−^ – G6P TSA complex (R=OH) there is a hydrogen bond between the 2-hydroxyl group of G6P and F_C_ of the MgF_3_^−^ moiety. This ^19^F resonance is significantly shifted upfield by 18.1 ppm in the βPGM-MgF_3_^−^-2-deoxy-G6P TSA complex (R=H) in H O buffer as there is no corresponding hydrogen bond present.

### Solvent induced isotope shifts (SIIS)

Solvent induced isotope shifts (SIIS) are defined as SIIS = δX(H_2_O) – δX(D_2_O) where δX(H_2_O) is the chemical shift of nucleus X in 100 % H_2_O buffer and δX(D_2_O) is the chemical shift in 100 % D_2_O buffer^38^. For free fluoride in solution, the SIIS observed is close to 3.0 ppm at 298 K, which represents a maximal effect and arises from through-space transmission of the electric field differences between F—H and F—D solvation. The extent of the isotope shift in enzyme-metallofluoride complexes depends on the number of the substitutable hydrogen atoms in the vicinity of the fluorine nuclei and also on the distance and angle of the coordinating hydrogen bonds^39,40^. For F···H–N and F···H–O hydrogen bonds between the metal fluoride moiety and the protein, the magnitude of the SIIS reflects the local proton densities around each fluorine nucleus^33,41^. A high proton density generates a deshielded fluorine nucleus resulting in large SIIS values and downfield chemical shifts approaching that of highly solvated free fluoride (Figure 5B, 5E). Conversely, the most upfield fluorine resonances always have the smallest SIIS values and line widths and are coordinated primarily by the essential catalytic magnesium cation with a concomitant low density of hydrogen bond donors. Partial deuteration can also be used to help the assignment of individual ^19^F resonances according to their coordination by Lys or Arg sidechains, and to determine the contributions to overall SIIS values by individual hydrogen atoms^16^.

### J coupling

The presence of well-defined and long-lived hydrogen bonds involving protein backbone amide groups and fluorine atoms in the active site has enabled the measurement of cross-hydrogen bond ^1^*J*_HF_ and ^2^*J*_NF_ scalar coupling constants in ^1^H,^15^N-HSQC experiments recorded without ^19^F decoupling. Here, the enzyme-metallofluoride complexes need to be prepared with ^15^N-labelled protein and the previously assigned backbone HN peaks are split horizontally by the one-bond ^1^H-^19^F coupling and vertically by the two-bond ^15^N-^19^F coupling for amide groups participating in hydrogen bonds to fluorine. Incorporation of ^19^F decoupling in both the ^15^N and ^1^H dimensions of the ^1^H,^15^N-HSQC pulse program results in a collapse of the scalar coupled peaks to an averaged position, confirming the involvement of fluorine in the effect (Figure 5D). The magnitudes of both coupling constants correlate closely with interatomic distances derived from high resolution crystal structures, providing independent validation of the similarity of enzyme-metallofluoride complexes in the solution and solid state (Figure 5C).

### NOEs

The close proximity and long residence times of fluorine atoms in near-transition state and ground state analogue complexes allows the specific assignment of fluorine nuclei to their atomic positions as defined in high resolution crystal structures. A driven nuclear Overhauser effect 2D ^191^H,^15^N-HSQC difference experiment results in different NOE profiles to nearby backbone amide groups, following selective irradiation of each of the fluorine resonances^20,21,42^. A transient 2D ^1^H,^19^F heteronuclear NOE experiment (HOESY) gives direct correlations between proximate ^1^H and ^19^F groups ^31^. Both NOE experiments require a residue specific backbone assignment for complete interpretation of the data.

### Dynamics

Generally for MgF_3_ ^−^ and BeF_3_ ^−^ enzyme-metallofluoride complexes, the fluorine nuclei are in slow exchange and give rise to discrete well resolved peaks in ^19^F spectra. Occasionally, particularly for AlF_4_^−^ enzyme-metallofluoride complexes involving nucleotides, the fluorine peaks are rotationally averaged, indicating that the active site architecture has a relatively low barrier to the rotation of the equatorial groups within the non-native octahedral geometry.

## Conclusions

So far, our results have identified no exception of analogue formation for wild type proteins that catalyse phosphoryl transfer requiring apical oxygen atoms that are either anionic or are hydrogen bonded to a catalytic base, including PSP, βPGM, PGK, cAPK, UMP/CMP kinase and RhoA/RhoGAP. This is reflected by the capability of forming the TSAs, especially MgF_3_ ^−^ complexes, which requires a more stringent setup in the active site of the enzyme than do fluoroaluminate complexes. Aluminium tetrafluoride has a much greater intrinsic stability in water, therefore TSAs containing it can be formed in wild type enzymes as well as in some of their mutated variants, as long as there is at least an anionic carboxylate as apical ligand. The trifluoroberyllate GSAs that have been tested only attach to an anionic oxygen from a carboxylate or a phosphate in all of these cases. Metal fluorides can be used to stabilize large complexes and study the intricate details of catalysis and are essential tools in the study of phosphoryl transfer enzymes.

## Accession Numbers

The coordinates and structure factors have been deposited in the Protein Data Bank under the following accession codes 2×13 for the PGK-ADP-3PG complex, 2×14 for the PGK(K219A)-AMP-PCP-3PG complex and 4axx for the PGK-ADP-BeF_3_^−^-3PG complex; 2wf6 for the βPGM-AlF_4_^−^-G6P TSA complex and 3zi4 for the βPGM-ScF_4_^−^-G6P TSA complex. Further information and requests for resources and reagents should be directed to and will be fulfilled by the lead contact, Matthew Bowler.

## Acknowledgements

This work used the platforms of the Grenoble Instruct-ERIC center (ISBG; UAR 3518 CNRS-CEA-UGA-EMBL) within the Grenoble Partnership for Structural Biology (PSB), supported by FRISBI (ANR-10-INBS-0005-02) and GRAL, and financed within the Université Grenoble Alpes graduate school (Écoles Universitaires de Recherche) CBH-EUR-GS (ANR-17-EURE-0003). We acknowledge the European Synchrotron Radiation Facility for the provision of synchrotron radiation facilities, and we would like to thank the staff of the ESRF and EMBL Grenoble for assistance and support in using beamlines ID14-2, ID29 and BM29. The BBSRC, UK and EMBL are thanked for financial support.

## Supplemental information

### NMR methods

All ^19^F NMR experiments were recorded at 298 K on a Bruker Avance 500 MHz spectrometer (operating at 470.38 MHz for ^19^F) equipped with a QXI (^1^H/^13^C/^15^N/^19^F) probe. Typically, ∼64k transients were acquired without ^1^H decoupling over a spectral width of 100 ppm and were processed using FELIX (Felix NMR, Inc., San Diego, CA) with backward linear prediction and sinebell functions shifted by 60°. All samples contained 1 mM β-PGM, 5 mM MgCl_2_, 10 mM NaF, 10 mM G6P and 2 mM NaN_3_ as a preservative, in HEPES buffer prepared in 100% D_2_O at pH 7.2 (corrected for D_2_O effects) except for the GaF_4_^−^ complex which was recorded at pH 6.0 (Ga^3+^ ions precipitate as Ga(OH)_3_ at higher pH values). The βPGM-AlF_4_^−−^-G6P complex was supplemented with 3 mM AlCl_3_, whereas the βPGM-ScF_4_^−^-G6P and the βPGM-GaF_4_^−^-G6P complexes were recorded using 10 mM of the appropriate metal chloride, which was necessary to out-compete the βPGM-MgF_3_^−^-G6P complex in both cases. Out of 10 metals screened we found that only scandium (III) and gallium (III) were able to form metal fluoride complexes with βPGM (Figure 3A).

## Crystallization methods

### PGK

Expression and purification of recombinant wild type HsPGK was as described^17,19^. Attempts to co-crystallise PGK in the closed conformation with ADP, AMPPCP, or BeF_3_^−^ failed. Therefore, crystals of the TSA complexes were subjected to soaks to obtain these complexes. Crystals of the PGK-3PG-MgF_3_^−^-ADP, the PGK-3PG-AlF_4_ ^−^-ADP and the PGK(K219A)-3PG-AlF_3_ ^−^-ADP complexes were obtained as described previously^17,19^, transferred to cryoprotection buffers (35% PEG 2000 MME; 0.1 M Bis/Tris pH 6.5, 20 mM DTT, 25 mM MgCl2 and 50 mM 3PG) lacking aluminium chloride or ammonium fluoride and containing 10 mM of the relevant nucleotide (ADP or AMP-PCP) and 0.1mM deferoxamine. For the PGK-3PG-BeF_3_^−^-ADP structure, the soak solution included 10mM ADP, 10mM NH_4_F and 10mM BeCl_2_. For the PGK-3PG-ADP complex, PGK-3PG-MgF_3_ ^−^-ADP crystals were soaked for 30 mins; for the PGK(K219A)-3PG-AlF_3_^−^-AMP-PCP complex, PGK(K219A)-3PG-AlF_3_^−^-ADP were soaked for 1 hour, and for the PGK-3PG-BeF_3_^−^-ADP, PGK-3PG-MgF_3_^−^-ADP crystals were soaked for 1 hour. Soaking HsPGK-3PG-MgF_3_ ^−^-ADP crystals or HsPGK-3PG-MgF ^−^-ADP with AMP-PCP was not successful. After soaking, the crystals were harvested, plunged into liquid nitrogen and stored at 100 K. Diffraction data were collected from cryo-cooled crystals to between 2.1 Å and 1.74 Å resolution on an ADSC Q210 CCD detector at beamline ID14-2 (λ=0.933 Å) at the European Synchrotron Radiation Facility (ESRF), Grenoble, France (see Table 1). Data were processed with MOSFLM^43^ and programs from the Collaborative Computational Project Number 4 (CCP4) suite^44^ or Phenix^45^. The structures were solved by molecular replacement using the HsPGK-3PG-MgF_3_^−^-ADP TSA complex (PDB accession code 2wzb^17^) for the closed conformation as search models with all ligands and water molecules removed. In all subsequent refinement steps, 5% of the data were excluded for calculating the free R-factor. Refinement was carried out alternately with REFMAC5^50^ or Phenix.refine^45^, and by manual rebuilding with the program COOT^46^. Ligands were not included until the final rounds of refinement so they could be built into unbiased difference Fourier maps. Stereochemistry was assessed with COOT with all residues in preferred or allowed regions.

### βPGM

For preparation of the βPGM-AlF_4_^−−^ –G6P TSA complex, 10 mM NH_4_F and 2 mM AlCl_3_ were added to the protein sample, followed by 5 mM G6P and the solution adjusted to a protein concentration of 15 mg ml^−1^. For crystallization, 2 μl of the AlF_4_^−^ inhibited protein was mixed 1:1 with 19-21 % (w/v) PEG 3350 and 50 mM Mg acetate and placed in sitting drop crystallisation plates. Large plate crystals appeared after 1 week and had the approximate dimensions 0.5 mm x 0.1 mm x 0.1 mm. For cryoprotection, the crystals were transferred to a buffer containing 22 % (w/v) PEG 3350, 50 mM Mg acetate, 2 mM AlCl_3_, 10 mM NH_4_F, 10 mM glucose-6-phosphate, 5 mM MgCl_2_, 50 mM HEPES pH 7.2 and 5% PEG 400 (v/v). Then the PEG 400 concentration was increased to 20% (v/v) in 5% steps (15 min at each concentration) by transferring the crystals between buffers with an increasing concentration of PEG 400. Crystals were harvested with a mounted LithoLoop (Molecular Dimensions Ltd., Newmarket, UK), plunged into liquid nitrogen and stored at 100 K. The PGM-ScF_4_^−^ –G6P TSA complex was prepared by adding 10 mM NH_4_F and 10 mM ScCl_3_ the protein sample, followed by 5 mM G6P and the solution adjusted to a protein concentration of 15 mg ml^−1^. For crystallization, 2 μl of the ScF_4_^−−^ inhibited protein was mixed 1:1 with the precipitant (26-30 % (w/v) PEG 4000, 200 mM sodium acetate and 100 mM Tris pH 7.5) and placed in sitting drop crystallisation plates. Small diamond shaped plate crystals appeared overnight with the approximate dimensions 0.01 mm x 0.01 mm x 0.001 mm. Crystals were harvested on a micromesh loop (MiTeGen, Ithaca, NY) and excess mother liquor removed before plunging into liquid nitrogen according to established protocols^47^. Diffraction data were collected from cryo-cooled crystals to 1.33 Å (βPGM-ScF_4_^−^-G6P TSA complex) and 1.3 Å (βPGM-AlF_4_ ^−^ –G6P TSA complex) resolution on an ADSC Q4R CCD detector on beamline ID14-2 (λ=0.933 Å) for native data and to 1.95 Å resolution on an ADSC Q315 detector on beamline ID29 (λ=1.77 Å, E = 7 keV) for the long wavelength data set of the βPGM-ScF_4_ ^−^ – G6P complex, at the European Synchrotron Radiation Facility (ESRF), Grenoble, France (see Table 2). Data were processed with MOSFLM^43^ and programs from the Collaborative Computational Project Number 4 (CCP4) suite^44^. The structure was determined by molecular replacement with MolRep^48^ using PDB accession code 2wf5^33^ as a search model with all ligands and water molecules removed. In all subsequent refinement steps, 5% of the data were excluded for calculating the free R-factor. Refinement was carried out alternately with REFMAC5^49^ and by manual rebuilding with the program COOT^46^.

## Figures

**Figure S1.**
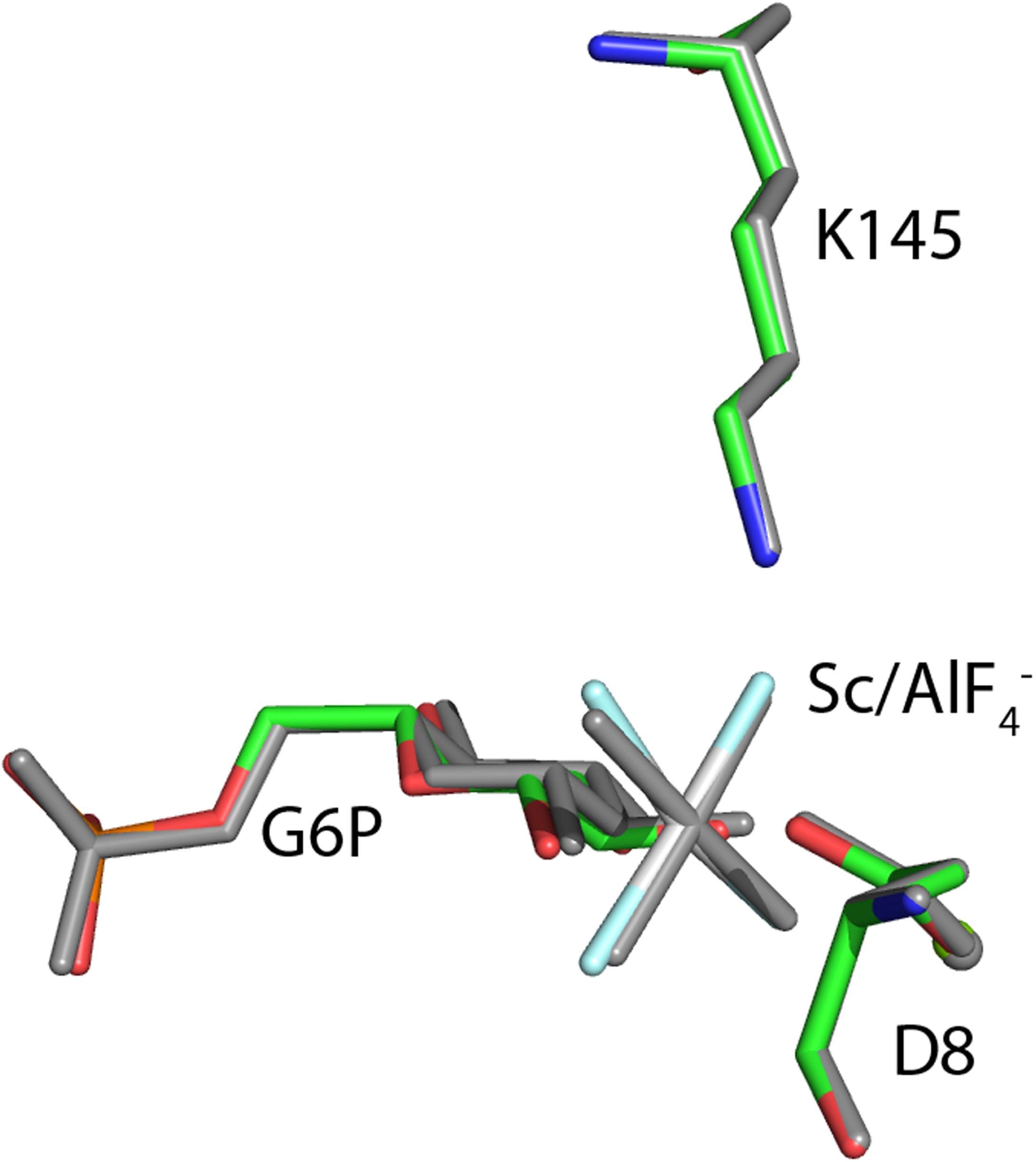
Comparison between the βPGM-AlF_4_^−^-G6P and βPGM-ScF_4_^−^-G6P TSA complexes. Carbon atoms are shown as green in the βPGM-ScF_4_^−^-G6P TSA structure (PDB 3ZI4) and all atoms are grey in the βPGM-AlF_4_^−^-G6P TSA structure (PDB 2WF6). The bond lengths are slightly longer in the ScF_4_^−^ moiety compared with the AlF_4_ ^−^ moiety (2.0 Å vs 1.8 Å), as the Mg-F distance is fixed this leads to a slight displacement of the central metal.

## Notes

### Competing Interest Statement

The authors have declared no competing interest.

